# Lost in the woods: shifting habitats can lead phylogeography astray

**DOI:** 10.1101/2024.07.03.601889

**Authors:** Richard A. Neher

## Abstract

Continuous phylogeographic inference is a popular method to estimate parameters of the dispersal process and to reconstruct the spatial location of ancestors of extant populations from samples viral genome sequences. However, these models typically ignore that replication and population growth are tightly coupled to spatial location: populations expand in areas with abundant susceptible hosts and contract in regions with limited resources. Here, I first investigate the sampling consistency of popular summary statistics of dispersal and show that estimators of “lineage velocities” are ill-defined. I then use simulations to investigate how local density regulation or shifting habitats perturb phylogeographic inference in continuous space and show that these can result in biased and overconfident estimates of ancestral locations and dispersal parameters. These, sometimes dramatic, distortions depend in complicated ways on the past dynamics of habitats and underlying population dynamics and dispersal processes. Consequently, the validity of continuous phylogeographic inferences is hard to assess and confidence can be much lower than suggested by the inferred posterior distributions, in particular when involving poorly sampled locations or extrapolations far into the past.

---

As organisms replicate, they also disperse in space, resulting in mixing of the population and exploration of new habitats. The reconstruction of ancestral locations, past migrations, and dispersal parameters from sampling locations of extant individuals, typically in combination with genome sequence information to infer the phylogeny, is known as phylogeography. Phylogeographic methods are implemented in popular evolutionary analysis software such as BEAST (Lemey *et al*., 2009, Lemey *et al*., 2010) and are widely used to study the spatial dynamics of viral populations (Dellicour *et al*., 2020). However, it has long been known that defining consistent models for demographic inference in continuous space is challenging (Felsenstein, 1975; Guindon et al., 2016).

In phylogeographic inference, one can either assume that individuals migrate between discrete locations or disperse in continuous space. Here, we consider migration in continuous space, where the underlying model is typically assumed to be diffusive, i.e. the probability of sampling a descendant at position **r**_*c*_ after time *t* when the parent was at position **r**_*p*_ is given by

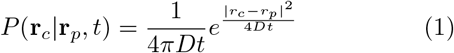

where *D* is a diffusivity with dimensions length^2^/time, we have assumed a two-dimensional space, and we have assumed dispersal is isotropic. Such a diffusive process is the simplest and most natural choice in absence of directed motion or long range migration. It is also mathematically convenient as unknown ancestral locations can be integrated in closed form and the marginal distribution of each position can be calculated exactly for a fixed tree along with the spatial likelihood. But this model assumes dispersal of one lineage is independent of other lineages, that dispersal properties are homogeneous across the habitat, and that the habitat does not change over time. Furthermore, it assumes that the tree topology and the branching rates of the tree are independent of the spatial location. Specifically, different components of the model – for example those describing the tree topology, coalescence properties, or molecular evolution – are independent of spatial likelihood and represented as multiplicative factors in posterior. A coalescent model, for example, would assume that each pair of lineage merges with equal rates, or a birth-death models would assume identical rates on contemporary branches of the tree. Variation of dispersal properties across the tree can be accommodated in relaxed random walk models that allow *D* in Eq. 1 to vary, but even in that case coupling between growth, location, and density is only be captured in indirect ways.

However, population dynamics are often tightly coupled to spatial location. Changes of the environment, for example, could mean that a habitat of a population is shifting, resulting in population growth and rapid branching at the expanding edge of the habitat and/or contraction in other areas. Similarly, pathogens spread in susceptible host populations that had not been previously exposed or have lost immunity. Examples of such shifting habitats are common and range from geological time scales, over decades (climate change), to months (seasonal fluctuations). Such changes are not restricted to the physical environment or population immunity, but ecological shifts can similarly affect the habitat. In all such cases, the replication process determining the phylogeny of the sample and the spatial location are strongly coupled. The extent to which violations of the independence assumption affect continuous phylogeographic inferences is not well understood.

Such coupling can in principle be accommodated in inference methods based on structured models that divide the population into discrete demes that could correspond to spatial locations that exchange migrants. Such models are flexible enough to allow for different population dynamics in different demes (Vaughan *et al*., 2014), though in practice can struggle to represent the population in sufficient detail or suffer from identifiability problems (Layan *et al*., 2023).

In a typical phylogeographic inference problem, the input data are genome sequences along with their sampling times and sampling locations. Software like BEAST will then sample the posterior distribution of the phylogenetic tree topology, the sampling times and locations of ancestral nodes in the tree, and model parameters like the diffusion constant *D* and the evolutionary rate *μ*. The inferred quantities of interest could be estimates of the ancestral location of a particular groups of samples, or summary statistics of the dispersal process. Here, I explore what dispersal quantities can be robustly estimated and how sensitive inferences are to the fact that coupling between population dynamics and spatial location is ignored. My focus here is only on estimates of dispersal parameters and ancestral locations and I will use simulations with perfect knowledge of the tree and the sampling locations. So no tree reconstruction will be necessary and sampling is uniform across the population. This simplification is justified since the tree topology tends to be informed much more strongly by sequence information. Using the true trees will result in clearer demonstrations of the limits of phylogeographic inference without the additional complication of tree uncertainty. Furthermore, I will assume strictly diffusive dispersal without any long-range migrations – the simulation and inference therefore use the same model. In practice, model violations through long-range migrations, errors in tree reconstruction, and sampling biases will add to the problems investigated here.

## Consistency of summary statistics of dispersal parameters

Before investigating the more complex issues of coupling between spatial locations and reproduction, I will first explore the properties of summary statistics in simple models where spatial location and population dynamics are independent. Popular summaries of the dispersal process are empirical estimates of the diffusion constant *D*_*w*_ (Pybus *et al*., 2012; Trovão *et al*., 2015) and a so-called *dispersal velocity v*_*w*_ (Lemey *et al*., 2010; Raghwani *et al*., 2011). These are calculated from the displacements along branches of the trees sampled from the posterior. For each branch *i* with parent *p*_*i*_, the displacement 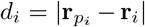 and the elapsed time Δ*t*_*i*_ are used to calculate

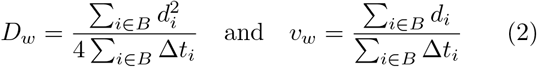

where *B* is the set of all branches of the tree. Effectively *D*_*w*_ divides the sum of all observed squared displacements by the total time elapsed on the tree to obtain an estimate of *D*. It is known as the ‘weighted’ estimate of the diffusion constant. The weighted dispersal velocity estimate compares the sum of absolute values of observed displacements to the total time, which has dimensions of a velocity. Alternative formulations use the mean of fractions 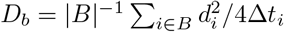 or *v*_*b*_ = |*B*|^−1^ ∑_*i*∈*B*_ *d*_*i*_*/*Δ*t*_*i*_ instead of the ratio of sums. We refer to these here as ‘by branch’ estimates. The latter tend to be dominated by short branches and are thus noisier (Trovão *et al*., 2015). The estimators in Eq. 2 are commonly used to summarize dispersal of lineages. While the estimator *D*_*w*_ directly estimates a parameter of the model using fundamental properties of diffusion (Pybus *et al*., 2012), it is unclear what *v*_*w*_ is measuring and how it behaves. I simulated diffusion along trees from Kingman coalescent (Kingman, 1982) and Yule model (Yule, 1925) tree ensembles and evaluated the different estimators using the known true ancestral locations. Kingman trees correspond to the classical neutral model of a constant size population and are characterized by many short branches and a few deep splits. Consequently, the spatial locations of samples from a Kingman tree tend to be clustered. Yule trees capture features of exponentially growing populations and tend to have a more even distribution of splits through time and long terminal branches. Examples for each ensemble are shown in Fig. 1.

**FIG. 1:**
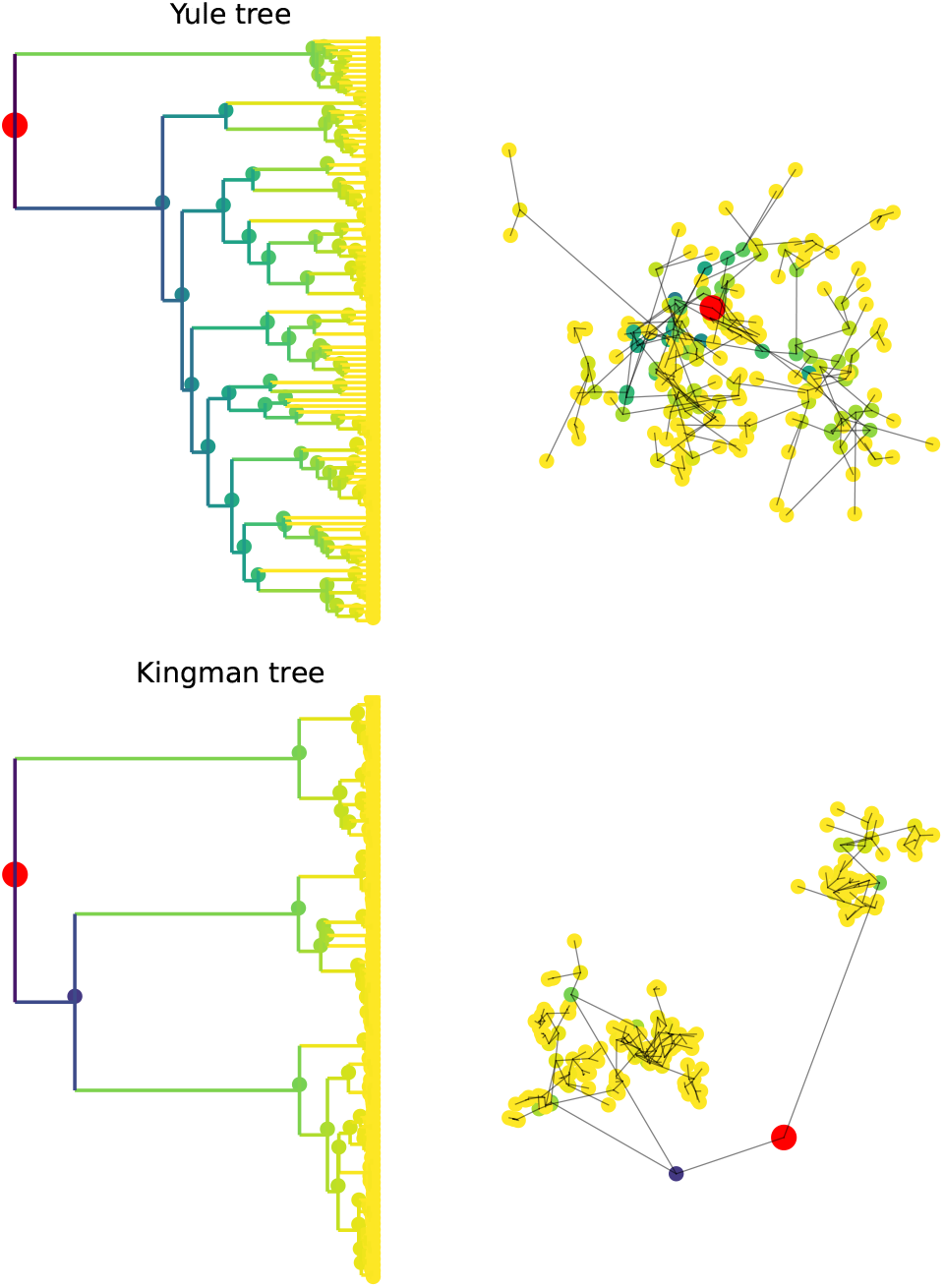
Illustration Yule and Kingman trees with *N* = 100 tips and the spatial location of the tips. The total tree length of Yule trees is proportional to the number of tips *n* and the time to the MRCA is log(*n*). Kingman trees have a total tree length that is proportional to log(*n*) and are characterized by rapid merging close to the present. Temporal and spatial scales are determined by parameter choice, only tree shape and spatial patterns are meaningful.

Fig. 2 shows empirical estimates of diffusion constants and velocity as defined in Eq. 2 from simulated data using freely diffusing locations along Yule and Kingman trees. The diffusion constant *D* can be estimated robustly from these data and the estimates are compatible with the diffusion constant *D* = 1 used to simulate the data. Values of *v*_*w*_ and *v*_*b*_ estimated from Kingman trees, however, depend strongly on the sample size and are incompatible with each other. That the velocity estimates are ill-defined is not surprising: By changing the sample size, the average length of branches in Kingman coalescent trees changes. Diffusion along a branch *i* over time Δ*t*_*i*_ results in a displacement of 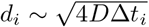 such that a quantity that depends on 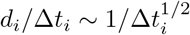 will change with typical length of branches Δ*t*_*i*_. With more samples in the tree, the average branch length decreases, and the velocity estimate increases. This expected scaling 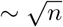 is indicated by the dashed line in right panel of Fig. 2. For Yule trees, the average branch length is independent of the number of samples and the velocity estimates are more stable. But as for the Kingman case, the estimates for *v*_*w*_ and *v*_*b*_ are not consistent with each other.

**FIG. 2:**
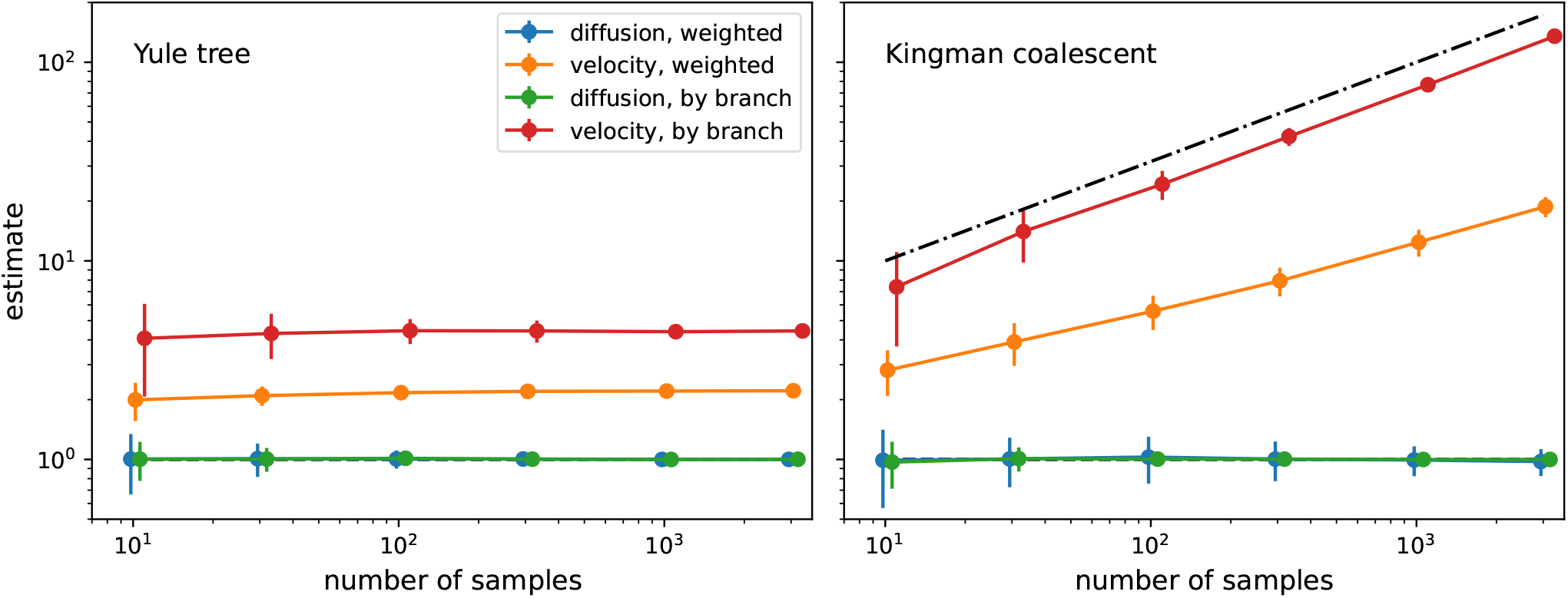
Empirical summary statistics of *D* and “dispersal velocity” for different sample sizes. While the diffusion constant *D* (equal to 1 in this example) can be robustly estimated from simulation data of the underlying model, the dispersal rate *v*_*w*_ (weighted) and *v*_*b*_ (by branch) are not well-defined quantities. For Kingman coalescent tree their numerical value depends strongly on sample size and there is no ground truth to compare to. The expected scaling 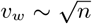 of *v* estimates is indicated by the dashed line.

In either case, estimates of *v*_*w*_ or *v*_*b*_ are not meaningful. The underlying model is diffusive and has no parameters that individually or combined would have dimensions of a velocity. There is no ground truth to compare the estimates to and *v*_*w*_ and *v*_*b*_ are just descriptive summaries of spatial spread that can not be compared between datasets. Similar observations have recently been reported by Dellicour et al. (2024)^1^.

While the estimator *D*_*w*_ as defined in Eq. 2 is well-behaved when the model is exactly diffusive, the estimator *D*_*w*_ is not necessarily well-behaved when dispersal occasionally proceeds via long-range migrations. If the probability of long jumps with distance *d* in the interval *x < d < x*+*dx* decays more slowly than ~ *dx/x*^3^, the expectation value of the *D*_*w*_ diverges. A popular model for long-range dispersal – as for example used in Dellicour et al. (2024) – are jumps drawn from a Cauchy distribution. The Cauchy distributions decays asymptotically as 1*/x*^2^ and does not have a finite mean nor variance. In this case the estimator *D*_*w*_ will be dominated by the longest jumps (or the size of the system) and fluctuate wildly^2^. When expectation of *D*_*w*_ diverges, the empirical value of *D*_*w*_ is no longer a useful summary of the typical dispersal process as it is determined by the most extreme events that fluctuate widely between different realizations of the process or even different samples from the same realization.

### Effects of density regulation on population structure and dispersal

Continuous phylogeographic inference typically assumes that the spatial diffusion process is independent of the branching pattern of the tree and ignores coupling between the two. Such a coupling, however, is expected since organisms compete for resources (e.g. susceptible hosts) and growth rates naturally depend on local population density. It has been known for a long time that ignoring such coupling leads to unrealistically heterogeneous population densities (Felsenstein, 1975), as is evident in the example of the Kingman tree in Fig. 1: The clades defined by long branches in the tree correspond to tight clusters of individuals in space. A more realistic population would spread out more evenly across the habitat.

If one simulates a diffusive spatial process in a square of length *L* with periodic boundary conditions, the population density shows strong fluctuations whenever the diffusion constant *D* ≪ *L*^2^*/T*_*c*_, where *T*_*c*_ is the coalescent timescale of a panmictic population (*T*_*c*_ = *N* in the case of a neutral model with a population of size *N*). I quantified this heterogeneity by binning individuals into two-dimensional bins with linear dimension *L/*5 (25 bins in total) and calculated the standard deviation of the number of individuals in these bins, normalized by the expected variation in a well-mixed population when each individual picks its location at random. The black line in Fig. 3A shows this heterogeneity measured by this standard deviation for a purely diffusive model as a function of the diffusion constant. For *D < L*^2^*/*2*T*_*c*_, diffusion is too slow to explore the entire habitat during the time since the MRCA, leading to clustering of individuals and high heterogeneity. This heterogeneity increases substantially for even lower *D*, when the population is essentially a cloud of width 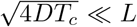. The comparison of *D* to the ratio *L*^2^*/T*_*c*_ will continue to be important below, and I therefore use diffusion constants in units of *L*^2^*/T*_*c*_.

**FIG. 3:**
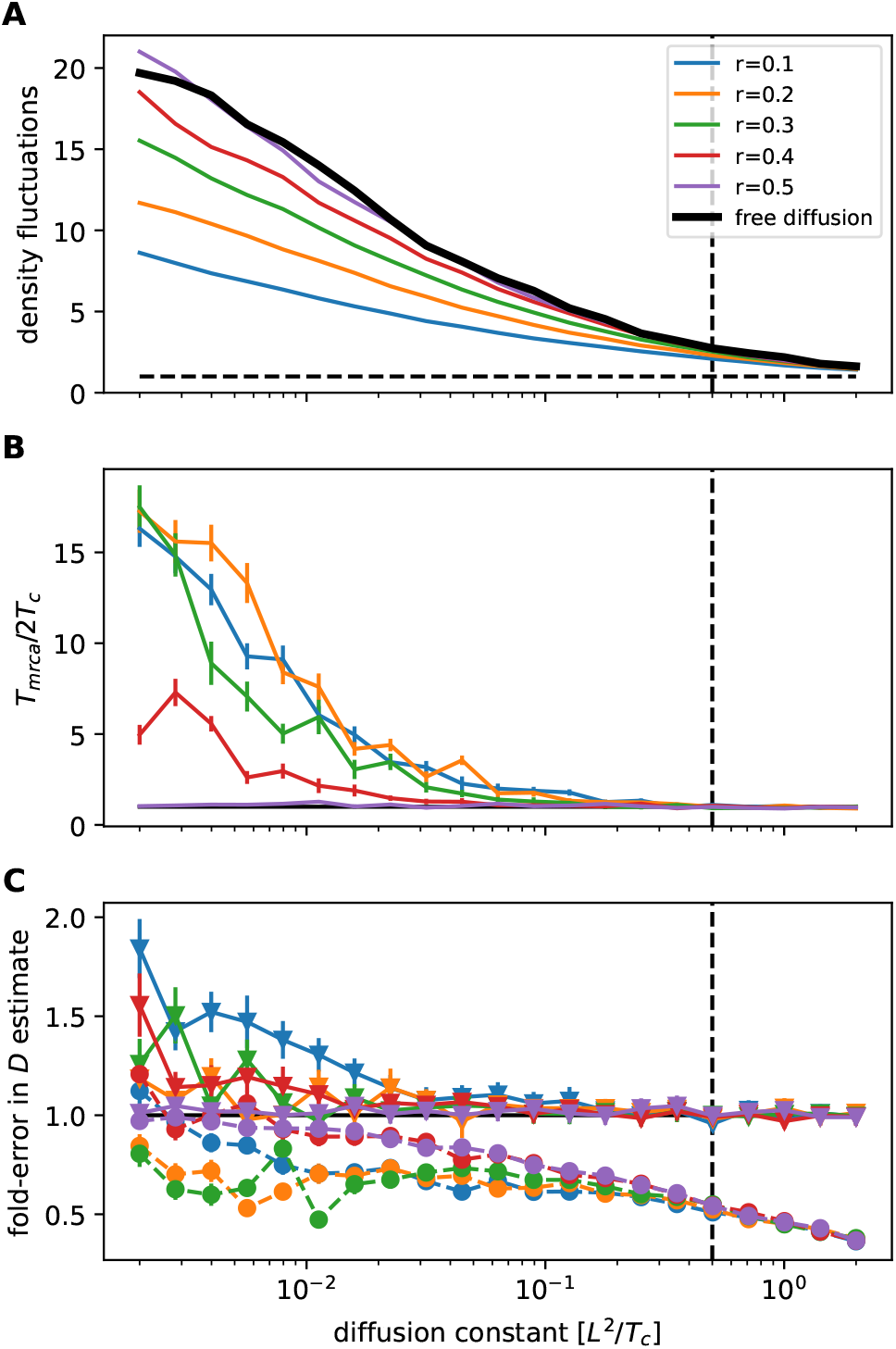
Effects of population density regulation. When spatial motion is independent of population density and coalescence is sufficiently rapid, the realized population density fluctuates strongly across the habitat. **Panel A** shows the standard deviation of the population density in bins of linear dimension *L/*5 as a function of the diffusion coefficient relative to the expected value if individuals were independently distributed across the habitat (dashed horizontal line). The thick black line shows the case of free diffusion, the colored lines show results from simulations with density regulation with different interaction radii *r*. For *r* ≪ *L*, density fluctuations are strongly suppressed. When density regulation homogenizes the population distribution, the time to the most recent common ancestor increases dramatically to values several fold higher than the neutral coalescence timescale *T*_*c*_ = *N* = 1000 (**panel B**), indicating that the population fragments in subpopulations that don’t interact. **Panel C** shows the fold-error of the dispersal estimator *D*_*w*_ for periodic boundary conditions (triangles, solid lines, like panels A&B) and reflecting boundaries (circles, dashed lines). The dashed vertical line indicates *D* = *L*^2^*/*2*T*_*c*_.

Biological populations spread into areas that support them and will quickly populate fertile areas that are below their carrying capacity. A simple model for the population density *ϕ*(**r**, *t*) would be

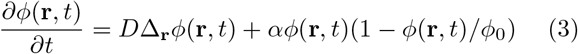

where Δ_**r**_ is the two-dimensional Laplace operator, *ϕ*_0_ is the carrying capacity, and *α* is the population growth rate at zero density. Such logistic density regulation was proposed by Bolker and Pacala (1997) and further analyzed by Etheridge (2004), but goes back to much earlier work by Fisher (1937) and Kolmogorov et al. (1937). The density has a stable fixed point at *ϕ* = *ϕ*_0_ we thus expect the population density to settle at a constant density *ϕ*_0_ and density fluctuations to decay with rate *α*.

While the dependence of growth (and thus tree structure) on density is a key component of real-world population dynamics, it is difficult to implement in phylogeographic inference frameworks. It is, however, fairly straightforward to include density dependence in forward simulations and these simulations can be used to investigate the effect of such coupling between growth and spatial location on phylogeographic inferences.

Instead of the coalescent trees used above, I used an individual based simulation forward in time. Individuals are assigned a fitness 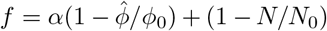 and produce a Poisson distributed number of offspring with mean 1 + *f*. The first term determines the fitness due to the local population density and has the tendency to stabilize this density at *ϕ*_0_, the second term (1 − *N/N*_0_) keeps the overall population close to the target population size *N*_0_. The population density is calculated using a Gaussian kernel with interaction radius *r*

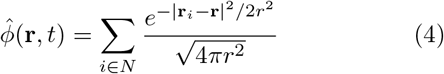

where the sum is over all individuals *i* in the population of size *N* and **r**_*i*_ is location of individual *i* at time *t*. The spatial locations of the offspring are chosen from a 2-D Gaussian centered on the parent with a variance to match the diffusion constant. The simulation records the tree of population, including their ancestral location and times. In addition to the true location of ancestral nodes, I also infer their locations from the locations of the extant samples. For this inference, I make the common assumption of continuous phylogeography that growth and location are uncoupled and that dispersal is diffusive. In this case, the marginal distributions of ancestral locations and their mean and variance can be calculated exactly using recursion relations.

The radius *r* of the density estimate 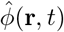 in Eq. 4 determines the distance over which different individuals compete for resources. As soon as *r* is much smaller than the habitat size, this density regulation leads to a more homogeneous population density, see Fig. 3A. But with decreasing diffusion, the population fragments into smaller and smaller subpopulations that interact only weakly and coalesce slowly. This manifests in an increasing time to the MRCA, see Fig. 3B, which at low *D* is limited by the time the population was simulated for. Spatial location and the coalescence process are thus strongly coupled.

Despite the substantial reduction in density fluctuations and strong population structure, the estimates *D*_*w*_ of the diffusion constant using Eq. 2 are not dramatically affected by density regulation, see Fig. 3C. If boundary conditions are periodic, that is the habitat has the topology of a torus, estimates *D*_*w*_ of the diffusion constant are accurate for most values of *D* but slightly inflated and noisy when the population is heavily fragmented (the increased noise is due to a smaller number of effective samples when tMRCA is large). If instead of periodic boundary conditions reflective boundary conditions are used, diffusion constants are underestimated once 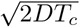 becomes comparable to the habitat size. This underestimation of diffusion constants will then result in over-confident inference of ancestral locations, in particular for recent ancestral nodes.

These simulations reveal that both at high and at low diffusion constants, assumptions of phylogeographic inference can be problematic: if diffusion constants are much smaller than *L*^2^*/T*_*c*_, the population is fragmented into effective subpopulations. This fragmentation violates the assumptions of phylodynamic models and in particular will affect estimates of population sizes. Such fragmentation could potentially be accounted for in structured coalescent model, but demes are not discrete and not known a priori such that these models are unlikely a practical alternative. If *D* ≫ *L*^2^*/T*_*c*_, the boundaries of the habitat restrict free diffusion, resulting in underestimated dispersal parameters, while estimates of deeper ancestral locations have uncertainties comparable to the habitat size and are thus uninformative.

### How do changing habitats affect phylogeographic inferences?

As discussed above, even a static carrying capacity in an equilibrium population can strongly affect population structure, though this structure has limited effects on estimates of ancestral locations and diffusion coefficients beyond boundary effects. In reality, carrying capacities will often vary through time. Such variation happens on all time scales, ranging from geological timescales over millions of years, environmental changes over millennia to seasonal fluctuations. The circulation of seasonal influenza viruses in temperate climates, for example, varies by several orders of magnitude between summer and winter.

Fluctuations in carrying capacity mean that populations not only spread because individuals move, but also because populations grow in regions where the population density is below the carrying capacity. In such situations, the accuracy and interpretation of phylogeographic inferences are unclear. In Fig. 4 we consider a case where the region with highest carrying capacity shifts from the left to the right in a periodic manner with period *T*. The carrying capacity is illustrated in Fig. 4A for 20 time points covering half a period. In between the two extreme left/right concentration of the carrying capacity (indicated by purple and yellow colors, respectively), the carrying capacity is uniform at a time between these two extremes. To avoid extinction at low dispersal rates, I implemented soft selection, meaning that the overall population size is kept approximately constant even if the entire population is stuck in the unfavorable region. If the dispersal rate is sufficiently high, however, the population follows the shifting carrying capacity and lineages in the tree undergo directed back-and-forth motion. Inference of ancestral locations in such situations can be misleading as illustrated in Fig. 4C. The population was sampled at a single time point when the carrying capacity is flat after having contracted to the left. The positions of older nodes and in particular of the root (symbols with bold red outline) are not estimated correctly. Their true locations are to the right of the current population, but the inferred locations are central in the shifted population. This is expected as there is no information in the current sample to suggest otherwise, but it highlights the potential of misleading inference, in particular when extrapolating far into the past.

**FIG. 4:**
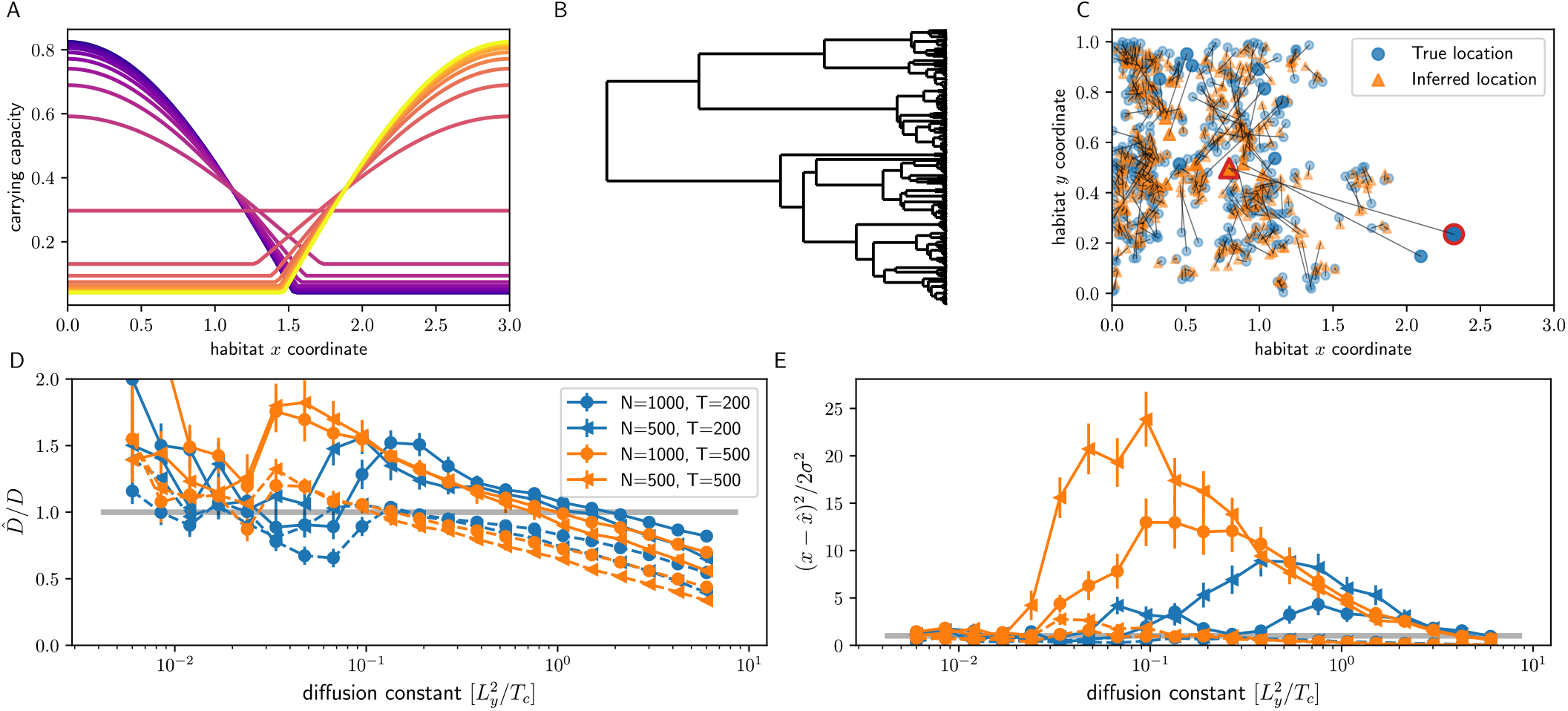
Phylogeographic estimates in rapidly changing environments. **Panel A** shows the time course of the carrying capacity through half a period. The color indicates time along the periodic habitat oscillations with yellow and purple representing most turning points of the oscillations. The carrying capacity first peaks on the left, becomes flat, and then peaks on the right before shifting back in the second half of the period. The habitat size is *L*_*x*_ = 3, *L*_*y*_ = 1. **Panel B** shows a tree from a simulation under these conditions with *T* = 500, *D* = 0.0008 and *N* = 500, while **panel C** on the right shows the inferred (triangles, orange) and true positions (circles, blue) of internal nodes, highlighting the root with large markers and a red border. The population is sampled at a time when the carrying capacity is flat after having moved to the left, that is the population is about to spread to the right. **Panel D** show estimates of diffusion coefficients in directions *x* (solid lines) and *y* (dashed lines) relative to their true values as a function of the true value of *D*. **Panel E** shows the average 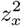 and 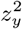 for the oldest 20% of internal nodes. Inference of the x-coordinate of ancestral nodes is drastically overconfident for some values of *D* (data for 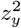 are hard to see in the figure as 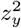 is of order 1 as expected). The summary inferences in panels D & E are made from many trees each of which represents samples from a single time point.

The severity of such biases depends on the period *T* at which the carrying capacity oscillates, on the population size, and on the dispersal rate. The bias tends to be strongest when the period *T* is of the same magnitude as the coalescent timescale. Fig. 4D shows the estimated diffusion constant (using Eq. 2) in *x* (solid lines) and *y* (dashed) direction relative to the true value used in the simulations. At very high diffusion constants, both *D*_*x*_ and *D*_*y*_ are underestimated due to the spatial constraints of carrying capacity as also discussed above (see Fig. 3). With decreasing *D, D*_*x*_ starts to be overestimated and peaks at a value that depends on the period *T*, before decreasing to (noisy) estimates that are broadly compatible with the true value. In this range of dispersal rates, the population can no longer follow the shifting carrying capacity and the *T*_*mrca*_ increases sharply (not shown). This decoupling happens at lower values of *D* for longer periods *T* of environmental change. With the parameters used in the simulation, the estimates of *D*_*x*_ deviate by a factor of 2 from its true value, which is not dramatic compared to the variation of dispersal rates in nature, but clearly shows that spatial inference can be substantially affected if the habitat changes on time scales comparable to the *T*_*mrca*_.

Along with problematic inference of diffusion coefficients, estimates of ancestral locations are over-confident. Panel Fig. 4E shows 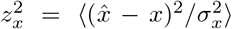 and 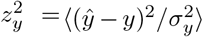 for the oldest 20% of internal nodes, where 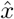 and *σ*_*x*_ are the mean and standard deviation of the posterior of *x*, respectively (and analogously for *y*). If the model estimated ancestral locations consistently, 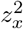 should be 1 on average. Instead, it is as high as 20 for the simulations shown and these over-confident and biased estimates depend in non-monotonic ways on the true diffusion constant and the period of environmental change. With decreasing *D*, 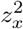 increases largely because the expected uncertainty decreases and the strongest over-confidence is observed when the dispersal capacity of the population is just about sufficient to follow the shifting carrying capacity. A peak value of 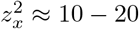 indicates that the typical ancestral location of the deep nodes is 3-5 standard deviations away from the true location, an example of which is shown in Fig. 4C. Conversely, at high dispersal rates, the average 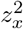 and 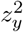 are both smaller than one, i.e. the inference is not confident enough. In this range, the estimated uncertainty is larger than the true uncertainty since the diffusion is constrained by the boundaries of the habitat while the estimated uncertainty extends beyond the habitat.

The above results show that behavior of a population in shifting habitats and the interpretation of phylogeographic inferences can depend strongly on the rate of environmental change and how it compares to the dispersal rate. At very low *D* the population can not follow shifting habitat and the situation is similar to the static case discussed above, which often leads to fragmented populations. With increasing *D*, the population starts following the shifting habitat, but inferred ancestral locations of deep nodes become unreliable. At high dispersal rates the expected uncertainties are larger than the habitat size and there is little information on deep ancestral locations.

To study the interplay of moving habitats and phylogeographic inference more systematically, I will now consider a situation where the population is constrained to a stripe with a Gaussian profile that moves at a constant velocity *v* along the *x*-axis (Fig. 5A). This set-up is sometimes referred to as “moving oasis” in the population genetics literature (Desai and Nelson, 2005). The dispersal rates necessary to keep up with the shifting carrying capacity can be estimated from the Fisher-KPP equation (Eq. 3) above. Eq. 3 admits solutions with a traveling front, i.e. the population “invades” the empty territory with velocity 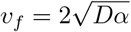 (Fisher, 1937; Hallatschek and Nelson, 2010; Kolmogorov *et al*., 1937). Note that the model itself does not have an explicit parameter that has dimensions of a velocity. The speed at which the population invades space is given as compound of diffusion and accelerated growth in regions of low density and only grows like the square root of diffusion constant.

**FIG. 5:**
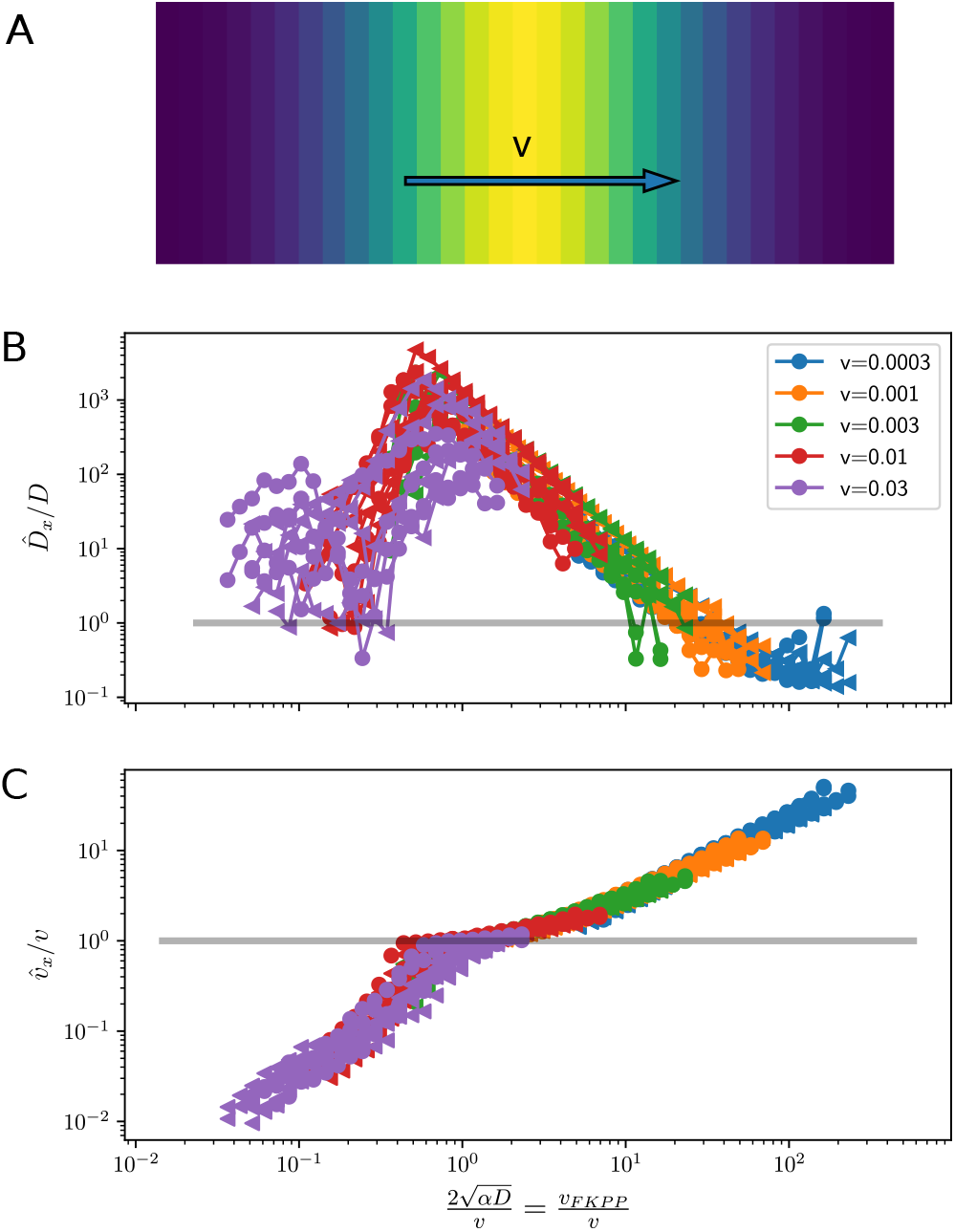
Phylogeographic inference in a moving habitat. **Panel A** shows an illustration of the simulated set-up: a carrying capacity with a Gaussian profile with *σ* = 0.5 that moves along the *x*-axis with velocity *v* on a patch with *L*_*x*_ = 3, *L*_*y*_ = 1, and periodic boundary conditions. **Panels B&C** show estimates of diffusion coefficients and velocities, relative to their true values, as a function of the ratio of the FKPP velocity to the external velocity *v*.

Fig 5B&C show simulation results for this traveling habitat. In Panel B, estimates of *D*_*x*_ using Eq. 2 are shown as a function of the ratio of the FKPP velocity *v*_*f*_ to the external velocity *v*. As expected, estimates of 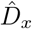 are often dramatically too high (by up to factors of 1000). They peak right around when the external velocity matches the FKPP velocity *v*_*f*_, corresponding to the situation where the population dynamics is maximally affected by the moving carrying capacity. At higher diffusion constants (to the right of the peak in Fig 5B), the effects of the moving habitat are less pronounced compared to the more vigorous undirected diffusion and the estimates 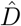 approach the true value as *D* increases. At lower diffusion constants (to the left of the peak) the population is unable to follow the moving carrying capacity.

I showed above that estimates of *v* using Eq. 2 are ill-defined in models with diffusive dispersal, but it is worth exploring how they behave in cases when the habitat is moving with an external velocity. Fig. 5C shows the estimator *v*_*w*_ in *x* direction relative to velocity of the habitat. This ratio is monotonically increasing with the diffusion coefficient and shows a range where it is close to 1, i.e. where the estimated velocity of lineages agrees with the external velocity.

These behaviors are explained comparing the speed *v*_*f*_ at which the population can invade new territory with the external speed *v*, the ratio of which is used as the *x*-axis in Fig. 5 B&C. For *v*_*f*_ ≪ *v*, the population cannot keep up with the moving carrying capacity and is saved from extinction only because selection is soft (Desai and Nelson, 2005). In this regime, the estimated velocity of lineages is below *v* and both the estimated *v* and *D* increase with *v*_*f*_. Once *v* ~ *v*_*f*_, the estimated lineage velocity coincides with *v*_*f*_, while the estimated *D* peaks at values orders of magnitude above the true value. Once *v*_*f*_ increases beyond the narrow range of *v* ≈ *v*_*f*_, the velocity estimate *v*_*w*_ increases further. In this latter regime, diffusive motions dominates over directed motion and the estimates of *v* suffer from the problems of sample size dependence and lack interpretation as shown in Fig. 2.

## Discussion

Phylogeography aims to reconstruct locations and migrations of ancestors from the spatial distribution of sampled individuals and their genome sequences. The latter allow the reconstruction of the phylogenetic tree which encodes how closely different individuals are related to each other. Together with the spatial location of the samples – the leaves of the tree – one can reconstruct the likely locations of ancestors and estimate parameters of dispersal. Popular tools for such inference, however, make strong assumptions about the dispersal process, including assuming that dispersal of individuals is diffusive (i.e. characterized by small displacements in random directions) and that the replication process is independent of spatial location. Here, I have explored the robustness of such inference to violations of the latter assumption while keeping microscopic dispersal strictly diffusive, i.e. I have avoided additional complications arising from long-range migration.

My first observation is that estimating dispersal “velocities” using displacements of lineages along the tree is problematic. Firstly, the numerical value of such estimates depends strongly on the sample size and tree ensemble. Secondly, the underlying model is diffusive and has no parameters that have dimensions of a velocity and even if the quantity could be estimated robustly, there is no interpretation of the quantity within the model. In cases where a population invades a novel habitat, e.g. the invasion of North America by the West-Nile virus (Pybus *et al*., 2012), the moving front of the population defines a bona fide velocity. However, as discussed above, this velocity is determined by the interplay of dispersal and population growth at the front, not a directional motion of individuals. It might be possible to capture this effective directional motions by random walks models that allow for directed motion (Gill *et al*., 2017). Estimators of a “lineage velocity” as defined in Eq. 2 only agree with invasion speed in a narrow, a priori unknown, parameter regime. In summary, the speed of invasion into a new habitat is better determined by modeling the position of the front (Pybus *et al*., 2012), rather than by tracking individual lineages along the tree.

A more complex set of questions concerns how coupling between growth rate of the population and spatial density affects phylogeographic inferences and how robust these inferences are to shifting habitats. As has long been known, ignoring the coupling between population growth and spatial location can lead to unrealistic population densities (Felsenstein, 1975) and that density regulation is necessary. However, once density regulation is in place, the population diversity and history depends strongly on whether dispersal is fast enough to explore the entire habitat during the time *T*_*c*_ it would take the population to coalesce in absence of spatial structure. If *D* ≪ *L*^2^*/T*_*c*_, the population fragments into subpopulations and the time to the most recent common ancestor increases dramatically compared to a neutral panmictic population. The resulting strong coupling between spatial location and coalescence process violates typical tree priors used in such inference and distorts estimates of ancestral population sizes. In the other case of *D* ≫ *L*^2^*/T*_*c*_, dispersal is rapid enough that uncertainties of ancestral locations of deep nodes are large compared to the size of the habitat and such inferences are thus uninformative. At the same time, dispersal parameters are likely underestimated for *D* ≫ *L*^2^*/T*_*c*_ since ancestral locations are constrained by the boundaries of the habitat. I did not consider additional effects that emerge from static carrying capacities that vary strongly in space, which can lead to unexpected effective lineage dynamics corresponding to sources where population densities are high and sinks where they are low (Hallatschek and Nelson, 2008; Wilkins and Wakeley, 2002).

Interpretation of phylogeographic inference becomes even more challenging when habitats shift in time. Depending on parameters, dispersal rates *D* can be overestimated or underestimated, sometimes by large factors, and estimates of ancestral locations can be overconfident or too uncertain. Simulations reveal that this behavior depends on the relative magnitude of the rate at which the habitat shifts and the FKPP velocity which depends on the diffusion constant *D* and the growth rate in empty regions *α*. The parameter range where both ancestral locations and dispersal parameters are inferred correctly is narrow and a priori unknown.

Phylogeographic methods are often studied using the spread of West Nile virus in North America and Rabies virus. West-Nile virus was first detected on the East Coast of the US in 1999 and reached the West Coast five years later in 2004 (Pybus *et al*., 2012), translating to a wave front velocity of about 1000 km/year. Pybus *et al*. (2012) estimated a diffusion coefficient between 200 and 5000 km^2^/day, where the latter refers to branches with particularly high diffusion constant. Dellicour *et al*. (2024) estimated a comparable diffusion constant of 310*km*^2^/day (via *D*_*w*_ rather than *D*_*b*_) for the expansion phase of the virus in North America and 2-3-fold lower rate after the initial expansion. These extremely high estimates of diffusive dispersal are implausible and likely indicate that directed elements of the spread of the virus are interpreted as undirected dispersal. Alternatively, one can interpret the spread from east to west as a range expansion. To this end, we need an estimate of the growth rate *α* of the population at low density, which we can roughly obtain from the fact that the virus is seasonal: over the course of a few months, its prevalence increases by orders of magnitude, which typically requires a doubling time of about a week or a growth rate of *α* = 0.1*/*day. Using 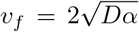 and equating it to 1000 km/year (≈ 3km/day), we obtain the estimate 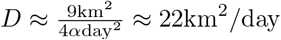 – around 10 to 200 times smaller than estimates from phylogeography. This lower estimate of *D* implies a daily exploration radius of about 5 km. This picture is also consistent with the apparent “slowing down” of lineages after the initial expansion across North America (Dellicour *et al*., 2024, 2020): Once the habitat was fully explored, directed range expansion with *v*_*f*_ ceases. Furthermore, rare long range dispersal could have strongly influenced the spread of the virus in ways not captured by Brownian or relaxed random walks (Hallatschek and Fisher, 2014). West Nile virus might for example by carried by migratory birds and their directed migration should not be described by a diffusion constant.

Diffusion constants of rabies virus are typically estimated to be between 500 and 1500 km^2^/year (Dellicour *et al*., 2017). The most recent common ancestor of various populations is between 30 and 100 years in the past. This would translate into rather limited spread of 150 to 500km of individual clades from their common ancestor, suggesting that these populations are highly structured (isolated by distance) and that their long range dispersal is dominated by rare introductions of the virus into new populations (Brunker *et al*., 2015).

The results presented here and the discussion of the examples above suggested that phylogeographic methods should be used and interpreted with caution. In many cases the habitat of a population has undergone dramatic and recurrent changes since the *MRCA* and such changes will affect inferences in ways that are not captured by the models. In addition, uneven sampling, or complete lack of samples from some regions, can undermine phylogeographic estimates (Kalkauskas *et al*., 2021; Layan et al., 2023). The ability to infer ancestral locations is further limited by long-range dispersal (Hallatschek and Fisher, 2014). To capture such deviations from simple diffusive motions, many inference models assume a “relaxed random walk”, where diffusion constant can vary across the tree according to a broad prior distribution (Dellicour *et al*., 2021). However, while such models will often generate a better fit, they don’t capture complex interactions between spatial location and population dynamics.

Why does phylogeography then often generate sensible results? On short time scales, a diffusive model will mostly reduce to a parsimonious reconstruction of ancestral locations which is robust to details of the model. Sampling at multiple time points further constrains ancestral locations sufficiently strongly that there is limited uncertainty, at least within the period for which samples are available. In these cases, phylogeographic reconstruction effectively interpolates between sampled locations along the relationship structure encoded in the tree and the results tend to be both plausible and well constrained by the data. But inferences for deeper nodes in the phylogeny long before the earliest samples can be problematic as the chance that the environment has changed and shifted, or that the true ancestral locations are in poorly sampled regions, increases. And while phylogeographic inferences will still often yield plausible results by virtue of their tendency to produce parsimonious solutions that aggregate available spatial information, they can also be highly unreliable and misleading. When population densities are not at equilibrium, there is no guarantee (and indeed we should not expect) that the posterior distributions suggested by such models accurately capture the uncertainty of ancestral events prior to the time interval covered by the data. While typical or most likely scenarios might often be predicted correctly, confidence statements should be treated with extreme caution. The tails of the inferred distributions are determined by properties of the model and the underlying assumptions typically do not reflect the wide spectrum of possible environmental histories. Model misspecification of the kind discussed here can strongly affect the probability mass in the tails of distributions. This concern particularly applies to scenarios population density, expansion, or contraction varies strongly in space and time. In such cases, model misspecification such as missing interaction between spatial location and population dynamics can lead to biased inferences.

A common approach in the field is to make models more complex and increase model flexibility by adding more parameters, for example by allowing for a broad distribution of diffusion constants in relaxed random walk models. Such expansion of parameter space and integration over additional phenomenological parameters like *D* will result in better fits and make the model more permissive to absorb misspecifications, but does not address the underlying problem. The computational resources necessary to infer such models increase rapidly with complexity, it becomes harder to understand how the data inform the inferences, and interpretation of inferred parameter values becomes ambiguous as they might have absorbed effects that were not explicitly modeled. In absence of a good understanding of the dispersal properties and the population history in time and space, simple approaches that allow transparent interpretation of how results are informed by features of the data are preferable to complex and computationally demanding models that can nevertheless fail to faithfully infer history or essential parameters.

## Materials and methods

All simulations were performed in Python and the associated code is available in the github repository github.com/neherlab/phylogeography.

## Acknowledgements

I am grateful to Louis de Plessis, Oskar Hallatscheck, Emma Hodcroft, Philippe Lemey, and particularly Simon Dellicour for stimulating discussions and useful feedback on earlier versions of this manuscript.

## Footnotes

1 I had alerted some of the authors of (Dellicour *et al*., 2024) in May 2023 to the problems with the velocity estimators. We independently explored the behavior of velocity estimators.

2 The largest jump in a sample of *n* jumps from a Cauchy distribution is typically on the order of 2*γn/π*, where *γ* is the scale of the Cauchy distribution. The second-largest jump is typically half, the third largest a third, etc. of the largest value. The estimator *D*_*w*_ sums their squares, which in many cases will be dominated by a few of the very largest values.

